# Stretch and Twist of HEAT Repeats Leads to Activation of DNA-PK Kinase

**DOI:** 10.1101/2020.10.19.346148

**Authors:** Xuemin Chen, Xiang Xu, Yun Chen, Joyce C. Cheung, Huaibin Wang, Jiansen Jiang, Natalia de Val, Tara Fox, Martin Gellert, Wei Yang

**Author notes:** These authors contributed equally (Xiang Xu, Yun Chen). Correspondence: Wei Yang, Martin Gellert.

## Abstract

Phosphatidylinositol 3-kinase-related kinases (PIKKs) are composed of conserved FAT and kinase domains (FATKIN) along with varied solenoid structures made of HEAT repeats. These kinases are activated in response to cellular stress signals, but the mechanisms governing activation and regulation remain unresolved. For DNA-dependent protein kinase (DNA-PK), all existing structures represent inactive states with resolution limited to 4.3 Å at best. Here we report the cryoEM structures of DNA-PKcs (catalytic subunit) bound to a DNA end, or complexed with Ku70/80 and DNA, in both inactive and activated forms at resolutions of 3.7 Å overall, and 3.2 Å for FATKIN. These structures reveal the sequential transition of DNA-PK from inactive to activated forms. Most notably, activation of the kinase involves previously unknown stretching and twisting within individual solenoid segments and coordinated shifts of neighboring segments in opposite directions. This unprecedented structural plasticity of helical repeats may be a general feature of HEAT-repeat proteins.

## Introduction

The DNA-dependent protein kinase (DNA-PK) is central to the process of non-homologous end joining (NHEJ) in both programmed gene arrangement and after unwanted DNA breakage (Davis et al., 2014). DNA-PK consists of a large catalytic subunit, known as DNA-PKcs, and the Ku70/80 heterodimer (Ku70/80) (Gottlieb and Jackson, 1993; Lees-Miller et al., 1990). DNA-PKcs is a Ser/Thr kinase of over 4000 residues and a member of the PI3K-related kinase (PIKK) family, which also includes mTOR, ATM, ATR and SMG1 (Baretic and Williams, 2014; Hartley et al., 1995). PIKK kinases play key roles in regulation of responses to nutrient stress (mTOR), misfolded and non-functional RNAs (SMG1), DNA double-strand breaks (DNA-PKcs and ATM) or single-stranded DNA (ATR) (Blackford and Jackson, 2017; Langer et al., 2020; Yang et al., 2017). The kinase activity of DNA-PKcs is modestly stimulated by DNA but becomes fully activated only in the presence of both DNA and Ku70/80, which is known to bind DNA ends (Chan and Lees-Miller, 1996; Hammarsten and Chu, 1998; Mimori et al., 1986; West et al., 1998). When activated, DNA-PK can phosphorylate both itself (auto-phosphorylation) and other repair factors (Meek et al., 2008). DNA-PKcs and Ku70/80 have also been implicated in telomere maintenance, RNA and ribosome biogenesis and the innate immune response to foreign DNAs (Hande, 2004; Meek, 2020; Shao et al., 2020).

Structures of the DNA-bound DNA-PK holoenzyme as well as DNA-PKcs associated with the C-terminal region (CTR) of Ku80 have been reported at 6.6 Å and 4.3-4.4 Å resolution, respectively (Baretic et al., 2019; Sharif et al., 2017; Sibanda et al., 2017; Yin et al., 2017). The first 3700 residues of DNA-PKcs are folded into ~65 a-helical HEAT repeats with a single β hairpin and arranged as an open N-HEAT (N for N-terminus) and a closed M-HEAT (M for middle) solenoid ring (Yin et al., 2017), which are nearly concentric, followed by the C-shaped FAT domain (Fig. 1a). The kinase domain at the C terminus occupies the hole in the FAT domain, and the two together, referred to as FATKIN (Baretic and Williams, 2014), form the “head” atop the double-ring “body”. The helical repeats are folded sequentially, going in opposite directions in the consecutive N- and M-HEAT rings. These two rings are linked back-to-back at the neck between the head and body but are splayed apart on the opposite side (the bottom), where DNA and the Ku subunits bind (Yin et al., 2017). Upon DNA and Ku binding, the first half of N-HEAT undergoes a rotational movement and converts the half-circle N-HEAT to a nearly closed ring (Yin et al., 2017) (Fig. 1, Video 1). The shape of DNA-PK and locations of domains are clearly defined, but at the resolution of 4.3 – 6.6 Å, functional interfaces between protein domains and between protein and DNA are less certain.

**Fig. 1.**
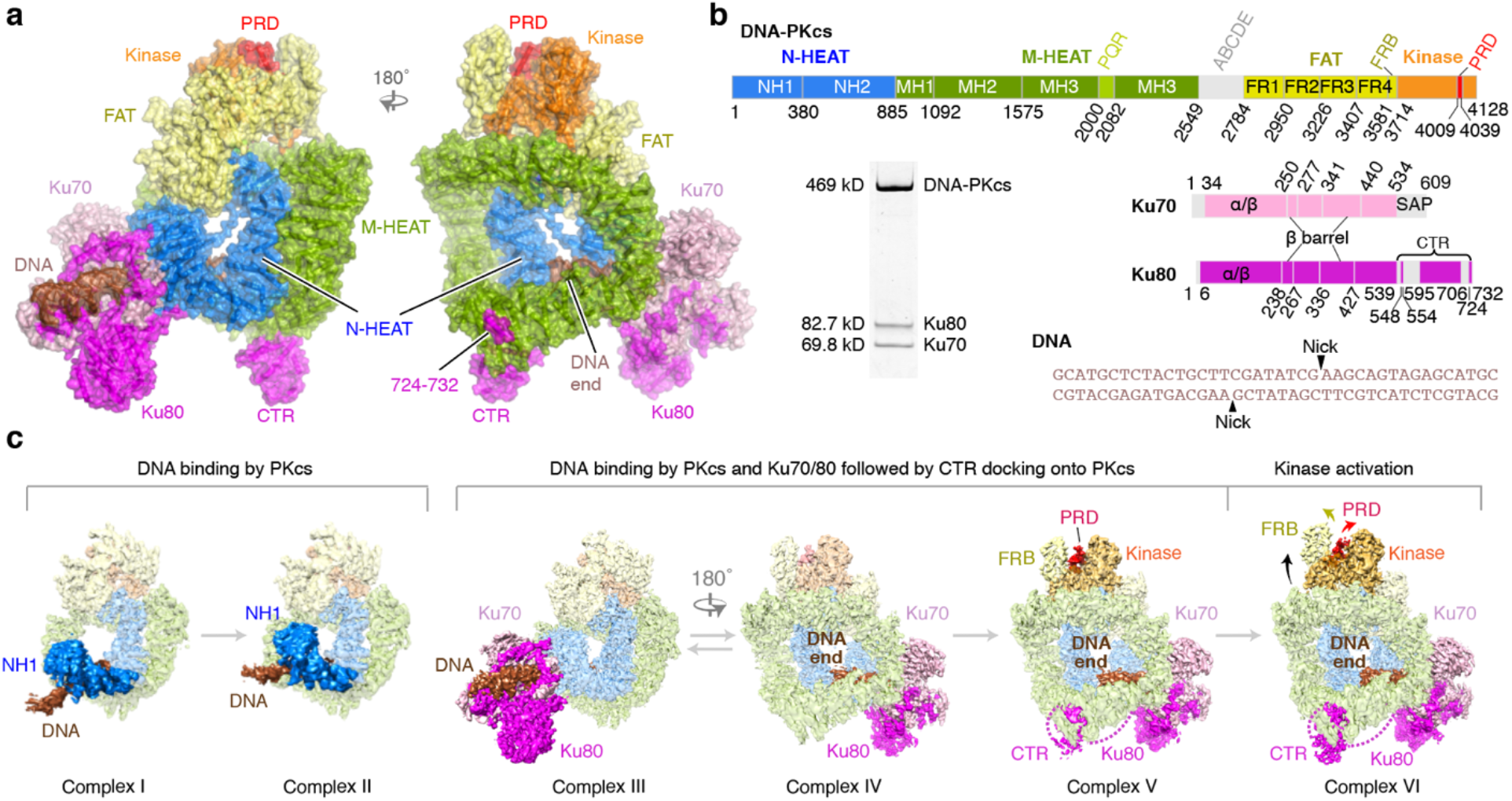
Structure characterization of DNA-PK complexed with DNA. **1a.** Front and back views of the holo DNA-PK complex VI. Major parts are labeled and color coded. **1b.** Diagrams of DNA-PKcs, Ku, and DNA sequences in the DNA-PK holo complexes. Quality of the protein sample is shown in the Coomassie-blue stained SDS gel. **1c.** The cryoEM density maps of DNA-PK complexes I – VI and how they are related. Regions that differ among these complexes are highlighted in colors shown in 1a, while regions unchanged are shown in muted colors. Complexes I-III are shown in the back view as the DNA changes the most among them, and complexes IV-VI are shown in the front view to illustrate the changes in Ku80-CTR binding and the activation of the kinase.

Multiple structures of other PIKK kinases complexed with accessory subunits, activation cofactors, ATP analogs, and even a substrate peptide have been reported, representing either active mTOR and SMG1 (Langer et al., 2020; Yang et al., 2017) or inhibited ATM and ATR (Jansma et al., 2020; Williams et al., 2020). In all cases the role of the HEAT-repeat structures, which is 5 to 9-fold larger than the kinase domain, remains unclear. Among PIKKs, DNA-PK is unique in its autonomous full activation by a short DNA duplex and auto-phosphorylation of its three subunits.

Here we report eight cryoEM structures of DNA-PKcs complexed with DNA or with DNA and Ku70/80 at 3.2 – 4.3 Å resolutions. These structures reveal the previously unknown function of DNA-end binding by DNA-PKcs, and in addition, an activated state of DNA-PK in association with the DNA end. Binding of Ku and a DNA end drive the intrinsic plasticity of DNA-PKcs toward kinase activation with coordinated stretching and twisting of helical repeats. The Slinky-like movement observed with DNA-PKcs may be general among HEAT-repeat proteins.

## Results

### cryoEM structures of DNA-PK

DNA-PKcs purified from HeLa cell nuclear extract was mixed with Ku70/80 and a 40 bp DNA that contained identical blunt ends with two internal nicks (Fig. 1b). Formation of the complete DNA-PK complex was confirmed by size-exclusion chromatography. CryoEM analysis of DNA-PK (see Materials and Methods) revealed three major species of protein-DNA complexes: DNA-PKcs-DNA, and inactive and activated holo-DNA-PK complexes (with Ku), two of which had more than one variant (Figs. 1c, S1). The FATKIN domains of the inactive and activated DNA-PK complexes were locally refined to 3.2 – 3.3 Å resolution (Table S1).

The two DNA-PKcs-DNA complex structures (I and II) at 4.3 Å resolution showed that DNA-PKcs alone can bind a DNA broken end and contact 15 bp. Complex I represented an initial DNA-binding state with DNA at the periphery of DNA-PKcs, whose protein structure was similar to a DNA-free DNA-PKcs (Sharif et al., 2017) (Fig. S2a), which was isolated from a sample containing DNA and Ku by cryoEM (Sharif et al., 2017). The structure of complex II showed that DNA and the associated N-HEAT region moved into the DNA-binding groove between N- and M-HEAT. The next three complexes (III to V) had DNA-bound to DNA-PK at 4.1 to 3.9 Å resolutions, with the Ku component increasingly resolved extending from the core (III-IV) to Ku80-CTR (V) (Fig. 1c). Structural differences between complexes III and IV probably reflected the dynamic flexibility of DNA-bound DNA-PK. With Ku80-CTR firmly docked, the structure of complex V became more stable. Interactions between DNA-PKcs and DNA were similar to those in complexes I and II, but specific contacts to the DNA end were formed in complexes III-V. While the protein and DNA components in complexes III-V and the previously reported DNA-PK structure (PDB: 5Y3R)(Yin et al., 2017) were similar, our complexes appeared to be expanded (Fig. S2b) and revealed specific binding of the DNA end and KU80-CTR by DNA-PKcs.

The activated DNA-PK complex (VI, at 3.7 Å resolution) exhibited extensive conformational changes. Compared to inactive complexes, the FATKIN head was raised, and the N- and C-lobes of kinase domain became open (Fig. 1c). Moreover, the PIKK regulatory domain (PRD, aa 4009-4039), which is partially disordered in mTOR and SMG1 (Langer et al., 2020; Yang et al., 2017), and closed in ATM, ATR (Jansma et al., 2020; Williams et al., 2020), and in all other DNA-PKcs and DNA-PK complexes, rotated 115° to expose the substrate binding groove. Complex VI also revealed that the activated DNA-PKcs has a potential binding site for inositol hexaphosphate (IP6), which has been shown to activate DNA-PK and NHEJ (Hanakahi et al., 2000). An IP6-binding site was first found in SMG1 and was predicted to be general among PIKKs, but had been absent in the existing DNA-PKcs structures (Gat et al., 2019), all of which represent inactive states.

### Structural comparison with apo DNA-PKcs

As our cryoEM density maps were of moderate resolution, to improve the structures and also for comparison, we took advantage of methionine locations well defined by selenium replacement of sulfur in the crystal structure of DNA-PKcs (PDB: 5LUQ, 4.3 Å) (Sibanda et al., 2017). In the process of building cryoEM models, we were able to re-refine the DNA-PKcs crystal structure (Table S2) (Fig. S2c), resulting in reduced R and Rfree (each by ~10%), reduced clash score and B values, and increased favorable Ramachandran distributions (Table S2). Re-assignment of secondary structures in DNA-PKcs was necessary to optimize this refinement (Fig. S3). The largest change was a 130-aa shift in the middle of the M-HEAT ring and involved over 200 residues in the crystal structure of DNA-PKcs (PDB: 5LUQ) and the cryoEM structure of DNA-bound DNA-PK (PDB: 5Y3R).

Domain and subdomain boundaries were determined by comparison of apo, inactive, and activated DNA-PKcs structures using Difference Distance Matrix Plot (DDMP) (Richards and Kundrot, 1988) (Fig. S4). According to established conventions (Sibanda et al., 2017; Yin et al., 2017), the three regions of DNA-PKcs are: N-HEAT (1-872), M-HEAT (890-2580), and FATKIN (2800-4128) (Fig. 2a-c). N-HEAT is broken into two segments (NH1 and NH2), each containing 8 helical repeats. NH1 (aa 1-380) binds 14 bp of DNA and interacts with Ku, while NH2 (aa 381-872) caps the DNA end and directly contacts M-HEAT and FATKIN (Fig. 2a). M-HEAT contains 32 helical repeats and assumes a heart shape in three segments (Fig. 2b). MH1 (aa 890-1092) forms domain connections at the neck; MH2 (aa 1093-1575) contacts NH2 and the N lobe of kinase at right angles; the large MH3 (aa 1576-2580) binds Ku and DNA and closes the M-HEAT ring (Fig. 2b). Based on DDMP analyses, M-HEAT, particularly MH3, is changed least among DNA-PKcs structures (Fig. S4).

**Fig. 2.**
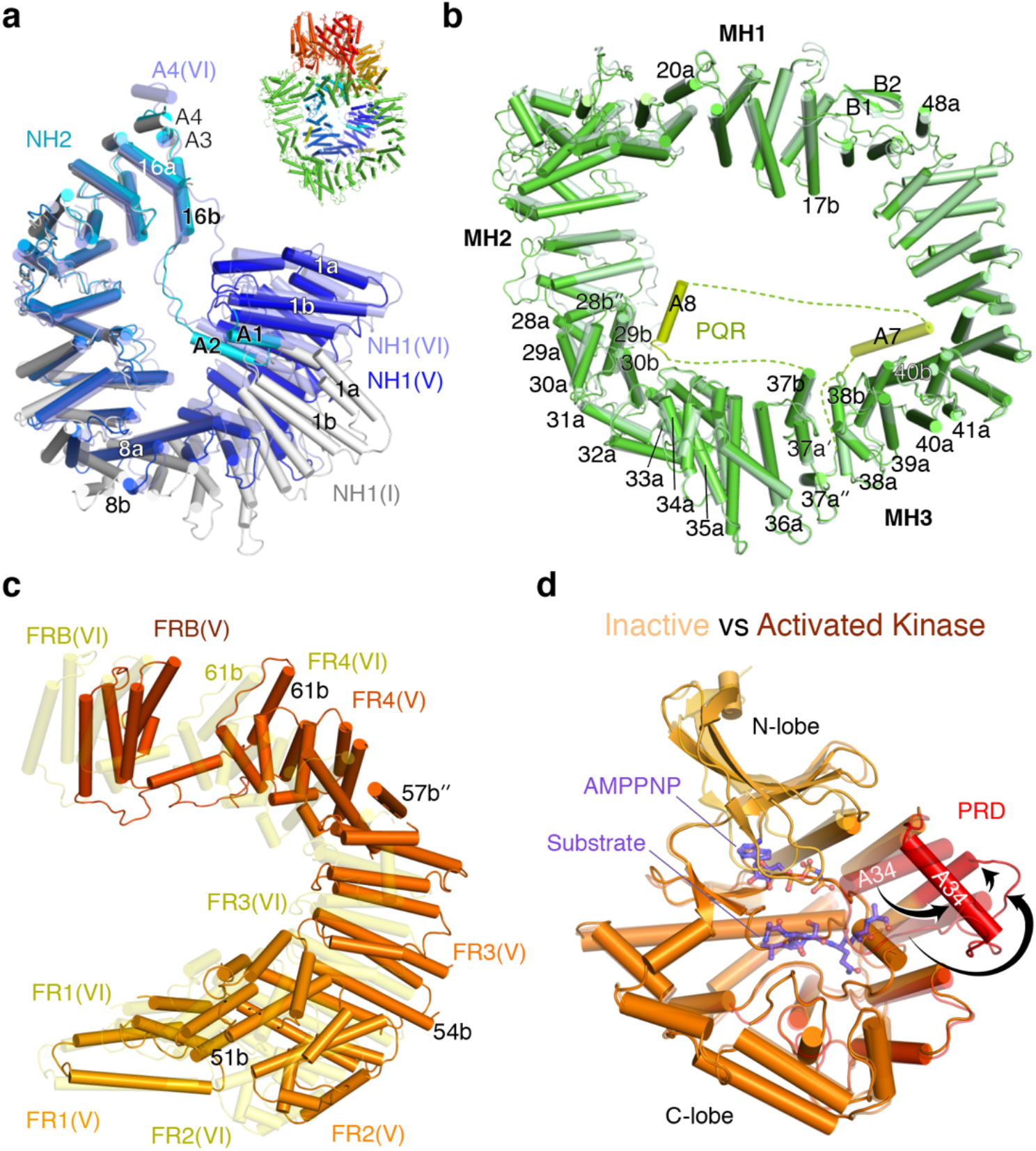
The DNA-PKcs structure. **2a.** The N-HEAT structures of Complex I (light grey), V (gradient colors of dark blue to cyan for residues 1 to 872), and VI (semi-transparent slate blue). The overall structure of DNA-PKcs is inserted as a reference. **2b.** M-HEAT structures in complexes V (solid green) and VI (semi-transparent pale green) are superimposed. MH2 (at the 10 o’clock position) is most changed between the two. Helices A7 (V) and A8 (VI) of PQR are highlighted in lime color. **2c**. FAT in complex V (solid orange to red) and VI (semi-transparent yellow) are superimposed at FR1. FAT is stretched between FR2 and FRB in complex VI (activated). **2d.** Kinase domain in complex V (semi-transparent) and VI (solid orange) are superimposed by the C lobe. AMPNPP and the substrate peptide (shown in semi-transparent purple sticks) are borrowed from the SMG1 structure (PDB: 6Z3R) after superimposing the conserved kinase domain. PRD in complex VI (highlighted in red) is open (as indicated by black arrows). The closed PRD in complex V would clash with the modeled substrate peptide.

The FAT domain starts at residue 2800, but the first segment FR1 (aa 2800-2944) actually moves with MH3 (Fig. 2c). The next segment FR2 (aa 2950-3199) contains non-repeating helices and interacts extensively with N- and M-HEAT at the neck, where the C lobe of kinase is also docked. The last section FR4 (3407-3564) interacts with the FRB (FKBP12-rapamycin-binding) domain and the N lobe of kinase, but FR3 (aa 3226 – 3394) between FR2 and FR4 has few interactions with the rest of DNA-PKcs. The conserved kinase domain (aa 3714 – 4128) (Fig. 2d) includes FATC (aa 4000-4128) as an intrinsic part of the C-lobe, as noted previously (Yang et al., 2013).

### DNA binding by DNA-PKcs

Previously, Ku80 was implicated in direct DNA-end binding by DNA-PK (Sibanda et al., 2017; Yin et al., 2017). In the better resolved cryoEM structures and crystal structure (Table S1, S2), the helices near the DNA end initially assigned to Ku80 were shown to belong to MH3 (as A7 in PQR, see explanation below) (Fig. 2b). Only DNA-PKcs has direct contacts with the DNA end and the proximal 15 bp. This observation is supported by observations that DNA-PKcs by itself has weak DNA-end binding activity (Cary et al., 1997; West et al., 1998) and that its kinase activity is slightly activated by DNA in the absence of Ku (Chan and Lees-Miller, 1996; Hammarsten and Chu, 1998).

In the two DNA-PKcs-DNA structures (I and II), only 24 bp of DNA are ordered (Fig. 3a). The 5^th^ to 15^th^ bp of DNA are bound by five helical repeats (3^rd^ to 7^th^, aa 161-331) of NH1 around the minor groove (Fig. 3b). Meanwhile the DNA end is contacted by the 9^th^ to 11^th^ helical repeats (aa 401-524) of NH2 (Fig. 3c). In converting from complex I to II, the DNA and NH1 move together by a rotation of 20°, which leads to a 10-Å shift from their being at the periphery of the M-HEAT ring to its center (Movie 2). As the DNA moves to the center, a pair of a helices (A1 and A2) (Fig. S3) extends from the 16^th^ repeat of N-HEAT toward it, with one side contacting DNA across the major groove and the other side contacting the 1^st^ and 2^nd^ repeats of N-HEAT (Fig. 2a, 3b).

Upon Ku binding and thus holo-DNA-PK formation (complexes III to VI), 37 of the 40 bp of DNA became stably ordered in these structures, as Ku bound DNA next to DNA-PKcs. In the transition from without (I-II) to with Ku (III-V), the entire DNA and the associated N-HEAT moved toward M-HEAT and also toward the outside of the M-HEAT ring by a 12° rotation and 8 Å translation, as if an arrow (DNA) were pulled (by Ku) further from the bow (M-HEAT) (Movie 3). In complexes III-V, the last helical repeat of NH1 (8^th^) finally makes contact with the DNA end, and in it N356, K357, and the positive helical dipole coordinate the 2^nd^ and 3^rd^ phosphates from the DNA 5’ end (Fig. 3c). The 5’ end, however, appears to be free. In contrast, the DNA 3’-OH is covered by W519, and the adjacent phosphates are bound by K518 and K520 (Fig. 3c). The asymmetric binding of 5’ and 3’ ends suggests that DNA-PKcs can easily accommodate a 5’ overhang (Fig. 3c). For the strand to continue as in DNA with a 3’ overhang or hairpin end, it has to bend away from the protein cap made of K518-K520.

The interactions of DNA with DNA-PKcs become more extended when the holo-DNA-PK complex shifts toward the activated state. As the groove between the N- and M-HEAT rings that sandwich the DNA becomes narrower, the interface between DNA-PKcs and DNA increases from 918 Å^2^ in complex V to 1036 Å^2^ in complex VI. Previously disordered loops (aa 801-817 and 839-846 of the 16^th^ helical repeat in N-HEAT) become ordered and contact M-HEAT at aa 2430-2511 (Fig. 3b). In addition to the protein-DNA interactions observed in complexes I-V, MH3 directly contacts the 4^th^ – 5^th^ bp via K2227, and the 9^th^ – 10^th^ bp via R2311 in complex VI (Fig. 3d). As the DNA is pushed against MH3 inside the HEAT rings, it is bent by ~20° away from M-HEAT and toward N-HEAT at the interface between DNA-PKcs and Ku.

### Ku stabilizes DNA-PKcs-DNA interaction

Ku is situated outside of DNA-PKcs and contacts the NH1 of N-HEAT and the MH3 of M-HEAT on either side of the DNA binding groove (Fig. 4a). Ku in the DNA-PK holoenzyme had essentially the same structure as in Ku-DNA complexes (Nemoz et al., 2018; Walker et al., 2001). When the Ku80 structures with and without DNA-PKcs were superimposed, the RMSD was 1.5 Å over 500 pairs of Ca atoms. The amino-terminal α/β domain of Ku70, which barely contacts DNA in the Ku-DNA complexes, binds MH3 (aa 2350 – 2420) and DNA (14^th^ – 16^th^ bp) tightly in complexes III-VI (Fig. 4b) by a rotation of 28° relative to the rest of Ku protein. The broad Ku70/80 base (composed of a/β and β barrel domains) becomes an extension of the MH3 of DNA-PKcs in binding DNA, protecting 15 bp (14^th^ to 28^th^) (Fig. 3d, 4a-b). While maintaining similar DNA interactions as Ku alone, the bridges of Ku70/80 contact the 3^rd^ to 5^th^ helical repeats of NH1 on the side opposite to the DNA (Fig. 4c). Therefore, Ku stabilizes the DNA-PKcs-DNA association by not only extending DNA binding, but also by fastening N-HEAT of DNA-PKcs onto DNA.

**Fig. 3.**
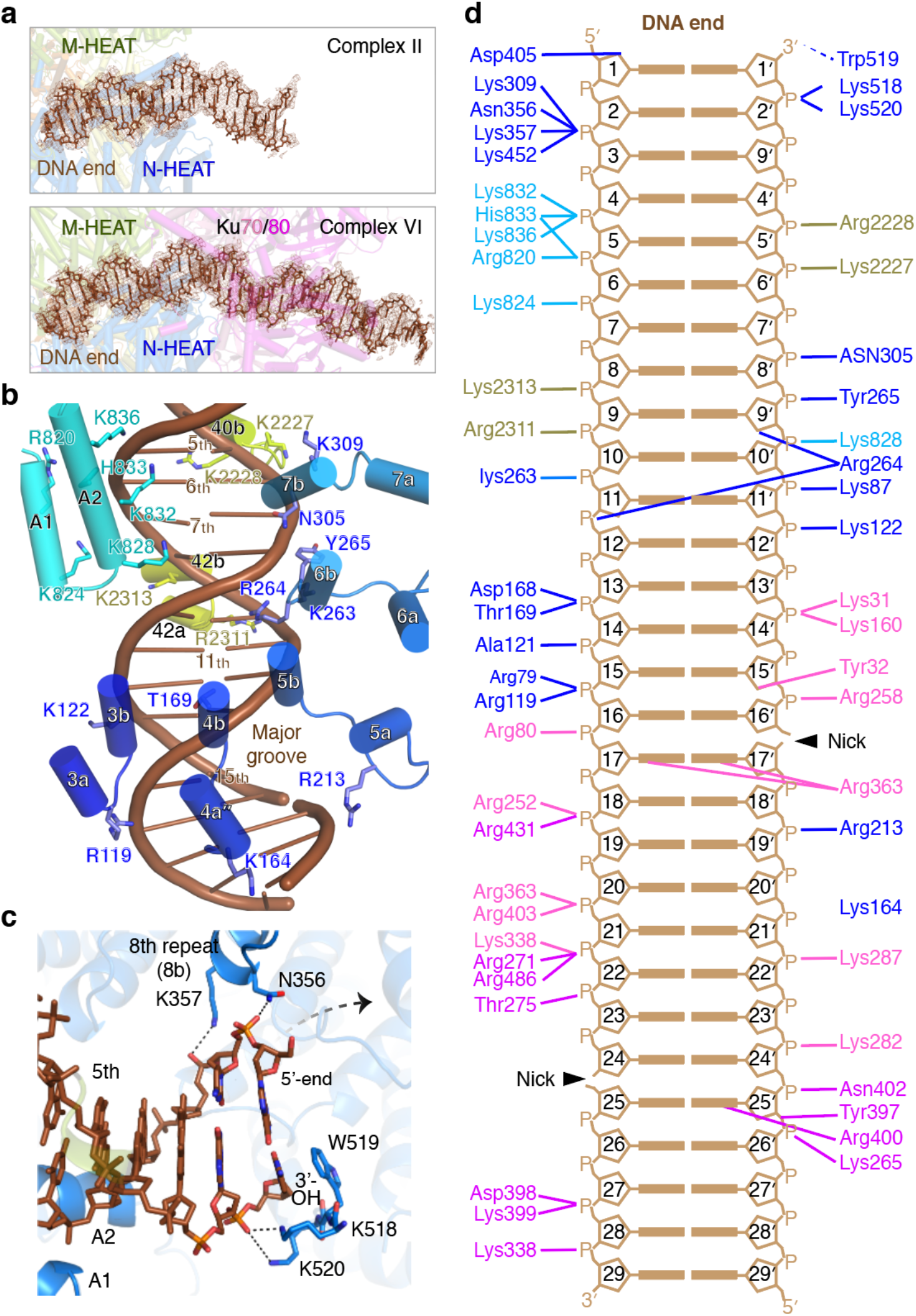
DNA binding by DNA-PK. **3a.** Representative density maps of the DNA in complexes II and VI were contoured color at 5.5 and 5.0 σ, respectively, in brown. Ku stabilizes the DNA in complexes III-VI. **3b.** Interactions between the DNA and NH1, the extension of the 16^th^ helical repeat (A1, A2) and MH3 in complexes II-VI. Helices A1 and A2 are not docked onto DNA in complex I. Residues of DNA-PKcs at the interface with DNA are shown as sticks and labeled. **3c.** In complexes III-VI, DNA-PKcs recognizes both DNA strands and caps the 3’ end specifically. Accommodation of a 5’-overhang is indicated by a dashed arrow. **3d.** Diagram of the complete DNA interactions made by DNA-PKcs and Ku in complex VI.

While the first 30 residues and the C-terminal SAP domain of Ku70 (aa 535-609) remained undetectable as in other Ku structures, Ku80-CTR (aa 542-732) becomes stable in complex V and VI structures (Fig. 1c). The folded domain in CTR (aa 595-706) is docked onto MH3 (aa 1700-1830), 65 Å away from the bulk of Ku, by a small interface of both hydrophobic and polar nature. In addition, a short helix of Ku80 (aa 548 – 555) is tucked between the 5^th^ and 6^th^ helical repeats of NH1 adjacent to the bridge and pillar of Ku70 (Fig. 4c). Selenium labeling of M731 (Sibanda et al., 2017) led to unequivocal identification of the last helix of Ku80 (aa 718-732), which is the only part of Ku80 present in the crystal structure of DNA-PKcs (Sibanda et al., 2017) (Fig. S2c-d). The last Ku80 helix is docked 20Å away at aa 1910-1970 of MH3, resulting in Ku contacting all four corners of the DNA-PKcs DNA-end binding groove in complexes V and VI (Fig. 4a).

**Fig. 4.**
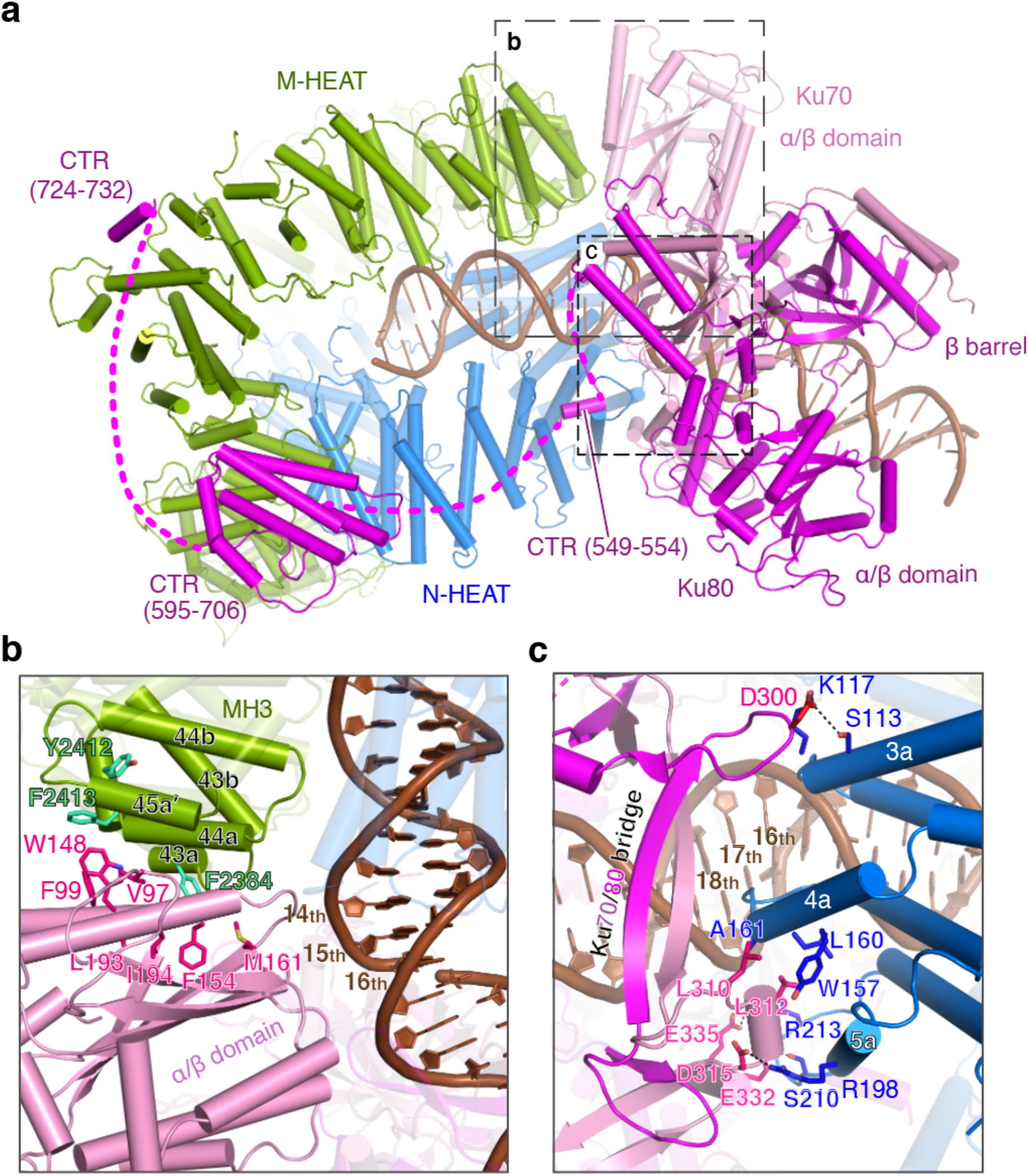
Ku association with DNA and DNA-PKcs. **4a.** A bottom view of the DNA-PK holo-complex (VI is used as an example). **4b**. The α/β domain of Ku70 interacts with DNA and MH3 of DNA-PKcs. Hydrophobic residues forming the interface are shown as sticks and labeled. **4c**. The bridges of Ku70/80, which binds the DNA major groove, interact with NH1 (3^rd^ to 5^th^ helical repeat), which interacts with the adjacent minor groove. The interacting residues are shown as sticks and labeled.

### Phosphorylation sites in DNA-PKcs

Residues 2581-2783 include the well-known auto-phosphorylation sites of DNA-PKcs, the ABCDE cluster (aa 2609-2647) (Ding et al., 2003; Neal et al., 2016). This 220-aa stretch was partially built in the apo crystal structure (PDB: 5LUQ and 7K17) inside the M-HEAT ring extending toward the center, where DNA would bind. In all six cryoEM structures, a relatively featureless density approximating the volume of the missing 200 residues was found outside of the M-HEAT ring and adjacent to the substrate binding groove of the kinase domain. We suspect that this density represents the mobile ABCDE region (Fig. S5) and speculate that the displacement of ABCDE from inside to outside the M-HEAT ring is correlated with DNA-end binding. Being near the kinase active site in the complexes with DNA, ABCDE may be phosphorylated in *cis* and facilitate downstream events in NHEJ (Crowe et al., 2020).

The second phosphorylation cluster (aa 2023-2056), known as PQR (Cui et al., 2005; Neal et al., 2016), is most peculiar. Among HEAT repeats of DNA-PKcs, PQR is the only part that does not follow a sequential folding pattern (Fig. 2b). As compounded by disordered linkers between helices, PQR caused the largest mis-tracing in the crystal structure (Sibanda et al., 2017). Instead of being near helix A7 (aa 2000-2018) and the 37^th^ helical repeat (Fig. S3), helix A8 (aa 2037-2045) of PQR is located 55 Å away and adjacent to the 32^nd^ and 33^rd^ helical repeats (aa 1575-1640) (Fig. 2b). The linker between A7 and A8 contains only 20 residues, which is a small number for the 55-60 Å separating the two. Interestingly, helices A7 and A8 co-exist in the crystal structure with A7 shifted 5 Å towards A8, but in each of our six cryoEM structures, either A7 or A8 is found, but not both. Helix A8 is dominant in four out of the five inactive complexes, whereas in the activated DNA-PK complex (VI) helix A7 is found in 75% of the population. Although disordered, the linkers inside of the M-HEAT ring have to cross paths with the DNA end and parts of N-HEAT. Although phosphorylation sites in the PQR cluster remain disordered, their locations suggest trans phosphorylation (as it is 90 Å away from the cis kinase domain) and potential influence on binding of DNA ends and repair partners inside the M-HEAT ring.

Other phosphorylation sites of DNA-PKcs (except for S3205) are in ordered regions, and their functions can be rationalized. S72 is adjacent to the bridges of Ku; its phosphorylation would interfere with interactions of DNA-PKcs and Ku and consequently destabilize DNA-PKcs-DNA binding. Indeed, phosphorylation of S72 has been reported to inactivate the kinase (Neal et al., 2011). T946 and S1003, whose phosphorylation has no effect on the kinase activity but inhibits the NHEJ pathway, may have additive effects with the neighboring ABCDE cluster in repair partner choices (Neal et al., 2011; Neal et al., 2016). T3950, which belongs to the activation loop of the kinase (Douglas et al., 2007), is buried in the available structures, and has to undergo conformational changes to be phosphorylated and modulate DNA-PK function.

### Activation of the kinase

In the activated state (complex VI), the position of FATKIN relative to N- and M-HEAT and its internal structure have changed dramatically (Movie 4) (Fig. 5a). In contrast, among complexes I to V, FATKIN moves little and only as a rigid body (Fig. S4, DDMP). The telltale sign of the activated state was the rotation of the PRD loop by 115° and opening of the binding groove for substrate (Fig. 2d). The positively charged putative IP6-binding site is nearby, wedged between FAT and kinase domains (Fig. 5b). Unlike mTOR and SMG1, when ATP or non-hydrolyzable ATP analogs were added to our sample, the DNA-PK complexes became highly heterogeneous and structural characterization was not possible. In the absence of ATP or ATP analogs, however, both the inactive and activated forms of DNA-PK were present in our samples.

**Fig. 5.**
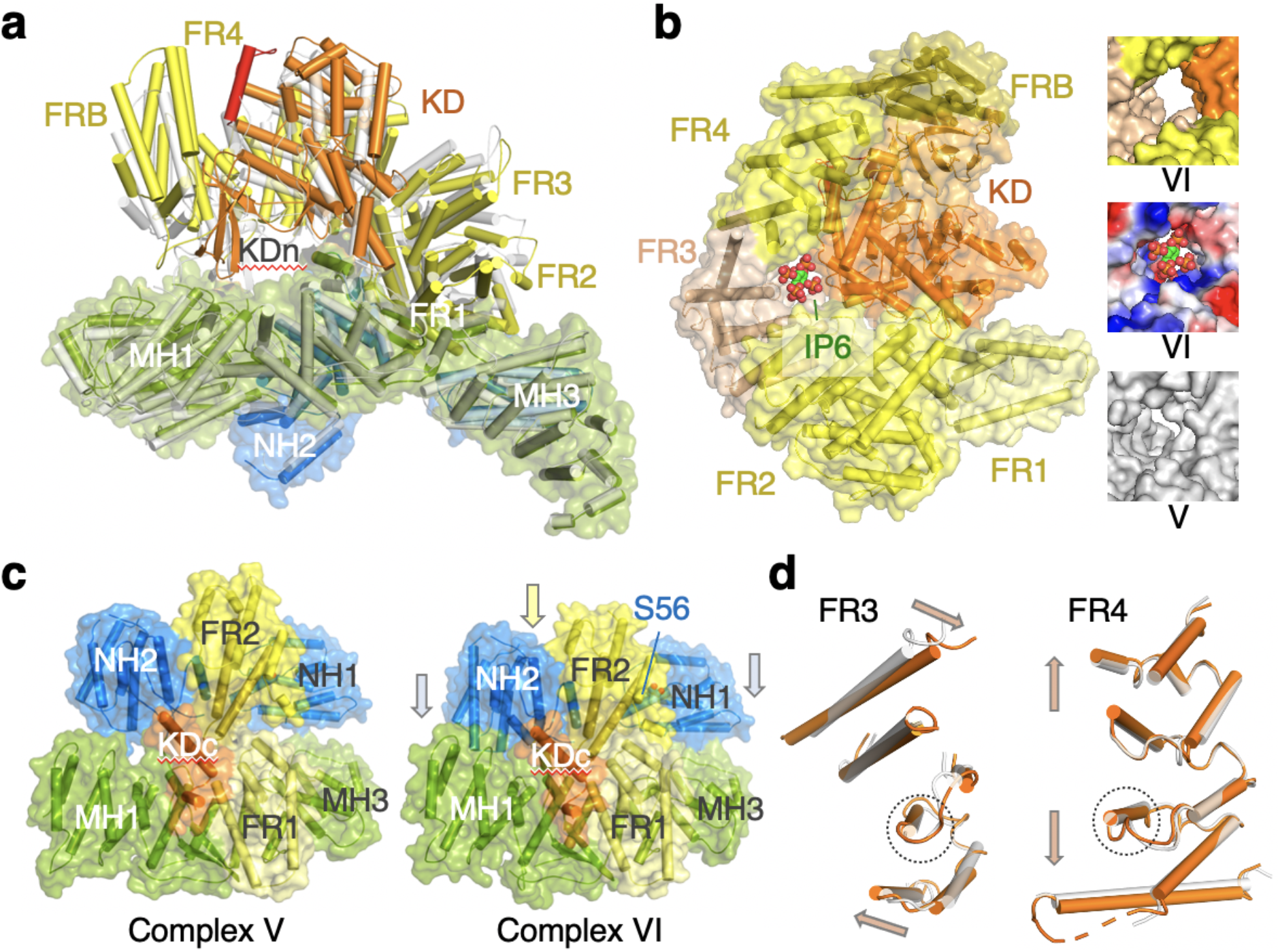
Kinase activation. **5a.** The FATKIN “head” in complex VI (multicolor) is raised relative to the body (covered molecular surface) compared with complex V (semi-transparent grey cartoon). **5b.** In complex VI, a positively charged pocket potentially binding IP6 is formed between the kinase domain (KD) and FR3. The pocket doesn’t exist in complex V as shown in the insert. **5c.** During transition from complex V and VI, the N-HEAT ring with FR2 moves toward the M-HEAT ring (MH1, MH3 and FR1), where the kinase domain (aa 3882-3916) is docked. S56 contacts FR2 only in the activated state (complex VI). **5d.** Comparison of FAT solenoid between complex V and VI. When one helix in FR3 or FR4 (in dashed circle) is superimposed, the surrounding helices twist (in FR3) or stretch (in FR4) in opposite directions as marked by arrowheads.

We asked how DNA and Ku activate the kinase activity of DNA-PKcs and what kinds of structural changes in the HEAT repeat rings are associated with this activation. Conformational changes from complex V to VI appear to emanate in the neck, where N- and M-HEAT, FAT and the C-lobe of the kinase converge. As shown in the animation (Movie 4), the NH1 and FR2 segments move toward each other to close the N-HEAT ring, and the entire ring including FR2 moves downward and enters the M-HEAT ring (Fig. 5c). Meanwhile, helix A4, linking N-HEAT to M-HEAT (Fig. S3), is squeezed out of the neck and rises up 4Å toward the kinase. Adjacent to helix A4, the C-lobe of kinase rises as well, together with FAT except for the FR2 segment, giving the impression of a raised FATKIN head (Fig. 5a). As the C-lobe is at the C-terminus of DNA-PKcs, but physically adjacent to FR2, the upward movement of kinase and downward movement of FR2 lead to unusual changes of every portion between them (Movies 4). As it is pulled in opposite directions, FR3 changes the twists between its helical repeats, and FR4 expands by increasing the distance between helical repeats (Fig. 5d). The solenoid made of FR3 and FR4 becomes more extended and more twisted in the activated state (Movie 5). As these changes happen, the PRD loop pops open. The importance of FR2 in kinase activation is supported by the observation that phosphorylation of S56 in NH1, which interfaces with FR2 in the activated state (Fig. 5c), causes kinase inactivation (Neal et al., 2011).

The dramatic changes in FATKIN are supported by changes in the two HEAT rings. As the N-HEAT ring enters, the M-HEAT ring expands (Movie 5), again by changing the distances and twisting angles between helical repeats. In the activated state, the N and C lobes of kinase also rotate away from each other, making room for substrate binding. This bidirectional movement appears to be correlated with the expansion of M-HEAT, which maintains interactions with the N-lobe of kinase by changing contacting residues.

DNA-PKcs alone is rather flexible, as evidenced in the differences between non-crystallographic symmetry mates (Sibanda et al., 2017) (Fig. S2d). But the intrinsic dynamics of DNA-PKcs are insufficient to re-arrange and activate the kinase. Binding of a DNA end leads to the re-organization of N-HEAT and brings NH1 and FR2 much closer than in apo structures, as shown in complex II. The closeness of NH1 and FR2 may explain the low kinase activity of DNA-PKcs in the presence of DNA (Chan and Lees-Miller, 1996). Binding of Ku70 and the entirety of Ku80 stabilizes the DNA-PKcs-DNA complex and shifts the ensemble of DNA-PKcs structures towards kinase activation.

## Discussion

The extensive global changes of DNA-PKcs that culminate to its activated state are results of local movements of helical repeats. Such movements within individual solenoids may be a common theme among HEAT-repeat proteins. For example, the fantastic shape changes of nuclear transport karyopherins exhibit local stretch and twist of helical repeats among large rigid-body conformational changes (Conti et al., 2006; Cook and Conti, 2010). Among PIKKs, binding of RHEB GTPase to mTORC1 also induces stretch and twist of helical repeats locally (Yang et al., 2017). In the case of DNA-PK, the extensive ensemble motions are supported by the binding of Ku and a DNA end. Although Ku is well known for its DNA end-binding activity, the cryoEM DNA-PK structures reported here reveal that Ku does not specifically bind DNA ends. The apparent DNA-end recognition is probably due to Ku’s closed topology, which requires it to bind DNA only via an open end by threading. Once on DNA, Ku has no direct contact with either the 3’ or 5’ end and can slide along following the DNA contour. This closed topology requires loading Ku first onto a broken DNA end in forming a functional DNA-PK holoenzyme. The association of Ku80’s last helix and DNA-PKcs is rather hydrophobic and DNA independent (Sibanda et al., 2017) and may enable the two to find DNA ends together. For DNA ends to be repaired, DNA-PKcs must yield the DNA ends to nuclease, DNA polymerase, other repair factors, and ligase IV. In the process, DNA-PKcs may slide inward along DNA with Ku. As shown in complexes V and VI, the full engagement of Ku with DNA and DNA-PKcs establishes a solid base of DNA-PK (Fig. 4a), on which extensive stretch and twist of HEAT repeats can take place and lead to the kinase activation. After seeing the extended and multifaceted stretch and twist of HEAT repeats in DNA-PK, we speculate that Slinky-like structural plasticity may be general among the vast number of HEAT proteins, PIKKs, karyopherins, condensins and cohesins alike. The trick for us is to catch them in action.

## STAR Methods

### Protein and DNA purification

DNA-PKcs was purified from HeLa cells (purchased from National Cell Culture Center, Minneapolis, MN) using a protocol we developed in 2009 based on a published method (Chan and Lees-Miller, 1996; Williams et al., 2008). The nuclear extracts were prepared according to the standard protocol (Abmayr et al., 2006) and then fractionated with 60% saturated ammonium sulfate. The precipitate was dissolved in DEAE loading buffer (50 mM HEPES pH 7.9, 75 mM KCl, 5% glycerol, 1 mM EDTA and 1 mM DTT) and clarified before loading onto a 200mL DEAE Sepharose FF column (GE Healthcare) preequilibrated with the loading buffer. DNA-PKcs fractions eluted from the DEAE column (in a gradient of 75-300 mM KCl) were pooled and diluted to a salt concentration of 150 mM KCl before loading onto a HiTrap Heparin HP column (GE Healthcare). A linear gradient of 150-500 mM KCl was applied to elute the protein, and the DNA-PKcs fractions were further purified using a Mono Q 10/100 GL anion exchange column (GE Healthcare) and eluted with a linear gradient of 150 to 350 mM KCl. The final purification step was on a Superose 6 10/300 GL size exclusion column (GE Healthcare) pre-equilibrated with 50 mM HEPES pH 7.9, 300 mM KCl, and 1 mM DTT. The purified DNA-PKcs protein was buffer exchanged to 50 mM HEPES pH 7.9, 300 mM KCl, 50% glycerol and 1 mM DTT, flash frozen in liquid nitrogen and stored at −80°C. All protein purification steps (including Ku purification described below) were carried out at 4°C, and protease inhibitors (100 mM PMSF, 1 mM pepstatin, 10 mg/mL aprotinin, 5 mg/mL leupeptin) were added to the buffer before each step of chromatography.

Ku70/80 protein was over-expressed and purified from HEK293T cells. Genes encoding full length Ku70 and Ku80 were cloned into pLEXm vector separately, and a His_6_-MBP tag and PreScission cleavage site were added to the N-terminus of Ku70 (Kim et al., 2015). HEK293T cells were pelleted 3 days after transfection and resuspended in 1/10 culture volume of lysis buffer (20 mM HEPES pH 7.9, 0.5M KCl, 5% glycerol, 0.5mM EDTA, 1 mM DTT and 1 tablet of Roche cocktail protease inhibitors). After sonication and centrifugation at 35,000 g for 1 h, the clear supernatant was applied to an amylose affinity column. After thorough wash, Ku protein was eluted in 20 mM HEPES pH 7.9, 0.5 M KCl, 5% glycerol, 40 mM maltose, 0.5 mM EDTA and 1 mM DTT. After removal of the N-terminal His_6_-MBP tag by PreScission Protease, the protein was loaded onto a Mono Q 10/100 GL anion exchange column (GE Healthcare) in 20 mM HEPES pH 7.9, 100 mM KCl, 1 mM DTT, 0.5 mM EDTA. Ku protein was eluted in a linear gradient of 100-500 mM KCl. The purified Ku70/80 fraction was buffer-exchanged into 20 mM HEPES pH 7.9, 150 mM KCl, 50% glycerol, 1 mM EDTA, 1 mM DTT, flash frozen in liquid nitrogen and stored at −80°C.

DNA oligos of 24 nt (5’-GCATGCTCTACTGCTTCGATATCG-3’) and 16 nt (5’-AAGCAGTAGAGCATGC-3’) were purchased from IDT (Integrated DNA Technologies, Coralville, IA) and purified using an 8-15% TBE-urea PAGE gel in small gel cassettes (Life Technologies). The oligonucleotides extracted from gel were loaded onto a Glen Gel-Pak column (Glen Research) and eluted with deionized H_2_O. These oligos were annealed in a buffer containing 20mM Tris-HCl (pH8.0), 50mM NaCl and 0.5mM EDTA in a Thermocycler to form a 40 bp DNA with identical blunt ends and two internal nicks (Fig. 1b). Such self-complementary DNA was initially designed as a cheap and easy way to make symmetric DNA of various length for co-crystallization with DNA-PK.

### Sample preparation and cryo-EM data collection

To assemble DNA-PK holoenzyme, purified DNA-PKcs, Ku70/80 and 40 bp DNA were mixed at the molar ratio of 1:1.2:1.2 in 50 mM HEPES pH 7.9, 100 mM KCl and 1mM DTT, and incubated at 4°C for 15 min. The mixture was purified over a Superose 6 10/300 GL column (GE Healthcare) pre-equilibrated with 50 mM HEPES 7.9, 100 mM KCl, 1mM DTT. Protein and DNA components of the holoenzyme were confirmed by SDS and TBE-urea PAGE gels. Fractions containing DNA-PK holoenzyme were pooled and concentrated to 0.5mg/ml for cryo-EM grids preparation.

The DNA-PK sample was loaded on either QUANTIFOIL R1.2/1.3 (Cu, 300 mesh) or Lacey grids (UC-A on holey 400 mesh Cu), 3 ul per grid at 100% humidity and 4°C in a Vitrobot, blotted for 4 s, and flash-frozen in liquid ethane. A total of 14742 micrographs from QUANTIFOIL grids were collected on the Titan Krios electron microscope operated at 300 kV at the Multi-Institute Cryo-EM Facility (MICEF) of NIH in the super-resolution mode of 130k nominal magnification (calibrated pixel size of 0.54 Å, corresponding to 1.08 Å at the sample level). An additional 7730 micrographs (4298 from QUANTIFOIL and 3432 from Lacey grids) were collected on a Titan Krios electron microscope operated at 300 kV at the Frederick National Laboratory (Frederick, MD) in the counting mode with a nominal magnification of 175K (calibrated pixel size of 0.86 Å). Lacey grids alleviated the preferred orientation problems manifested with QUANTIFOIL grids.

### Structure determination and model refinement

The software MotionCor2 (Zheng, 2017) was used for drift correction, during which dose-weighting was applied and the pixel size was binned to 1.16 Å/pixel to merge all micrographs from the two microscopes. CTF (contrast transfer function) estimation was measured with the dose-unweighted micrographs using Gctf (Zhang, 2016). 2,945,665 particles were picked on dose-weighted micrographs using Gautomatch (developed by K. Zhang; https://www.mrc-lmb.cam.ac.uk/kzhang/Gautomatch) and extracted with RELION-3.0.8 (Fernandez-Leiro, 2017) using a box size of 352 × 352 pixels. An initial map was obtained with cryoSPARC (Punjani, 2017), and two-dimensional (2D) projections were generated for template-based particle picking. The re-picked 5,592,709 particles were subjected to 2D classifications in RELION-3.0.8. After excluding 3,258,196 bad particles, three-dimension (3D) classifications in RELION and cryoSPARC with and without alignment were applied to classify different conformations. The resulting good maps and the associated particles from 3D classifications were selected for further classifications and refinements according to the standard procedure (Fig. S1). All reported resolutions were determined based on the “gold standard” of the 0.143 Fourier shell correlation (FSC) criterion (Swint-Kruse, 2005). Local resolution was estimated using ResMap (Kucukelbir et al., 2014). For model building, we used the published 4.3-Å resolution apo DNA-PKcs structure (PDB: 5LUQ) (Sibanda et al., 2017) as the initial model to build cryo-EM structures of the 3.2 Å inactive FATKIN and the 3.7 Å DNA-PK complex VI, which then were used as the template for building Complexes I-V. We first fit the coordinates into the cryo-EM maps using Chimera (Pettersen et al., 2004), and then manually adjusted and rebuilt the models according to the cryo-EM density in COOT (Emsley, 2010). Real-space refinement in Phenix (Adams et al., 2010) was used to refine the models, and MolProbity (Chen et al., 2010) was used to validate the final model. The refinement statistics are summarized in Table S1. The detailed classifications and map qualities of the 8 structures reported in this manuscript are shown in a supplemental figure (Fig. S1).

For comparison and validation of our cryoEM structures of DNA-PK, we also rebuilt and refined the DNA-PKcs crystal structure (Sibanda et al., 2017) using the diffraction data deposited with the Protein Data Bank. Using our 3.7 Å cryoEM DNA-PK complex VI structure as the initial model, we got a solution by Molecular Replacement and rebuilt the DNA-PKcs structure against the 2 Fo-Fc as well as Se anomalous maps in COOT. The model was iteratively refined using strategies of rigid body, group B factor, TLS parameters, XYZ (reciprocal-space and real-space), NCS application and secondary structure restraint in Phenix. MolProbity was used to validate the final model. A total of 7293 residues were built per asymmetric unit, which included 3629 and 3646 residues of DNA-PKcs in chain A and chain B, respectively, and the C-terminal residues 724-732 of Ku80 in chain C and chain D. Rwork and Rfree of the re-refined structure of DNA-PKcs complexed with the C-terminal helix of Ku80 were 0.29 and 0.34 at 4.3 Å, respectively. The detailed refinement statistics are summarized in Table S2.

## Data Availability

The structures and cryoEM maps have been deposited with EMDB and PDB with accession code of 7K19, 7K1B, 7K1J, 7K1K, 7K1N, 7K0Y, 7K11,7K10 and EMD-22622, EMD-22623, EMD-22624, EMD-22625, EMD-22626, EMD-22618, EMD-22620, EMD-22619. The re-refined crystal structure has been deposited in PDB with accession code of 7K17. These data will be released immediately upon publication.

## Author contributions

J.C.C. developed protocols for DNA-PKcs purification and complex assembly; X.X. and Y.C. improved DNA-PK complex preparation and collected cryoEM movies; X.C. and Y.C. processed cryoEM data, X.C. completed model building and structure analysis; H.W., N.d.V. and T.F. helped with cryoEM data collection, J.J. troubleshot cryoEM data collection and analysis, X.C., M.G. and W.Y. wrote the paper.

## Acknowledgements

We are grateful to Drs. K. Meek and D. Leahy for critical reading of the manuscript.

## Supplemental Information

**Table S1.**
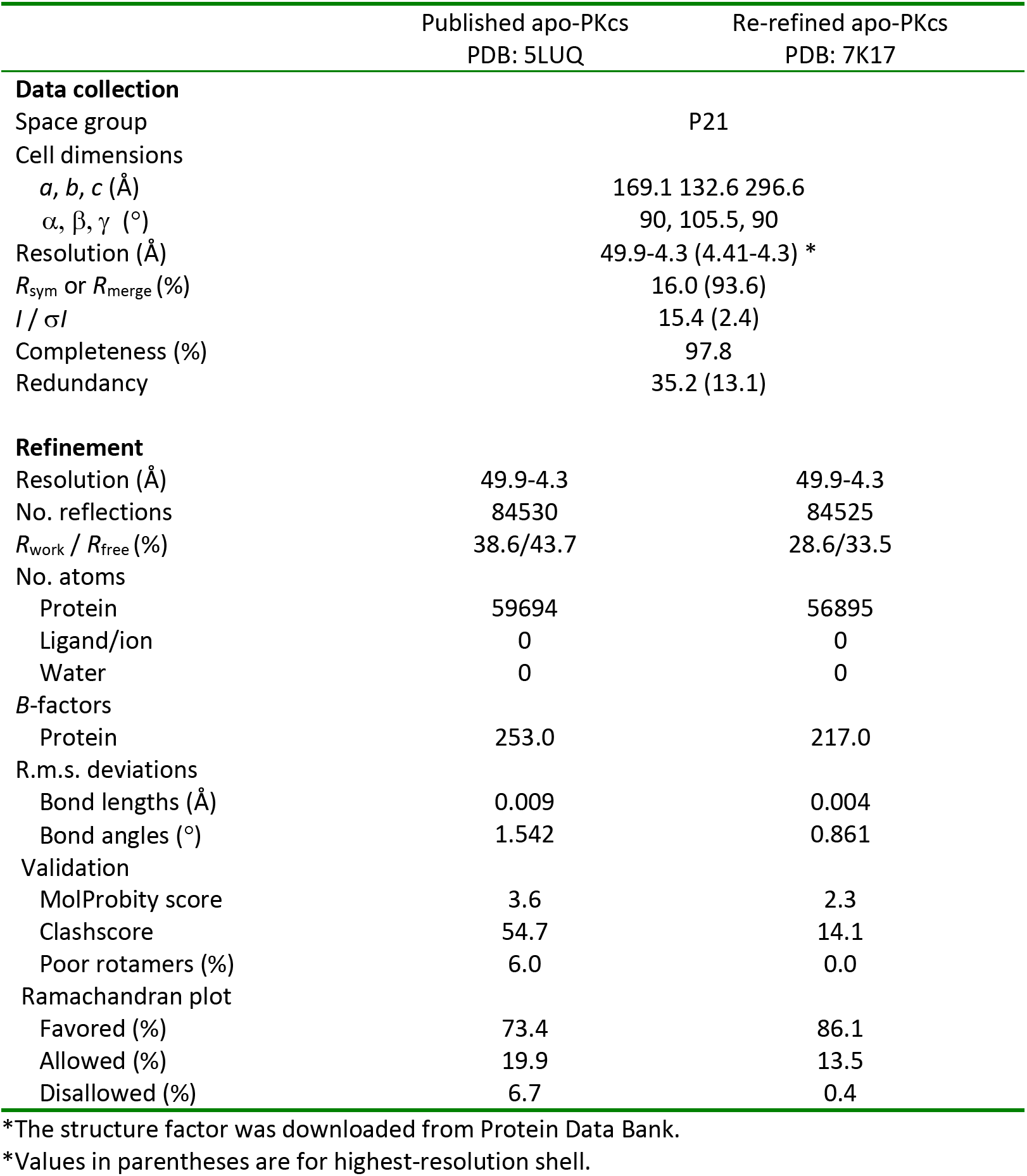
Statistics of re-refinement of the DNA-PKcs crystal structure.

**Table S2.**
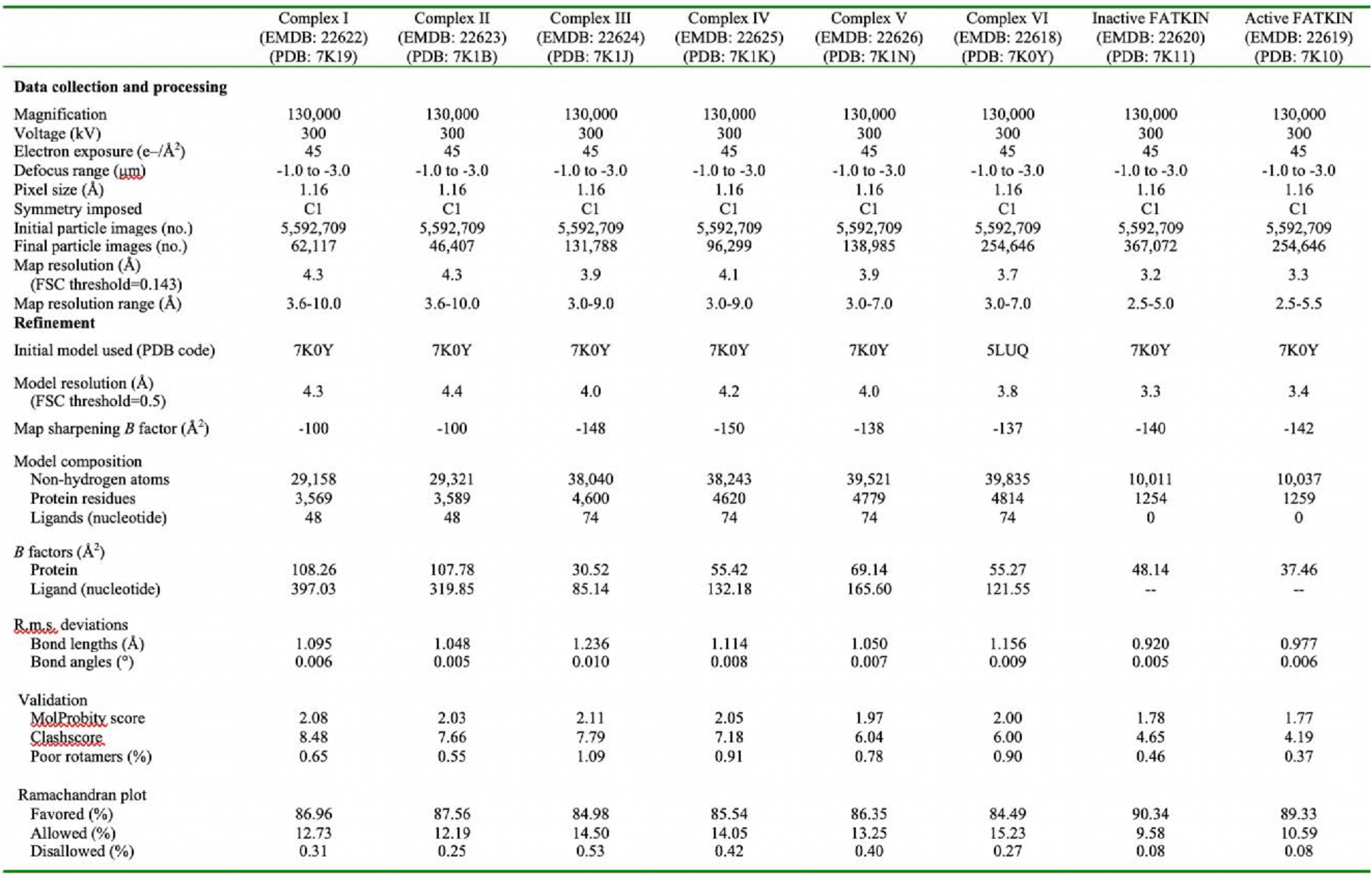
Statistics of cryo-EM data collection, refinement and validation.

**Fig. S1.**
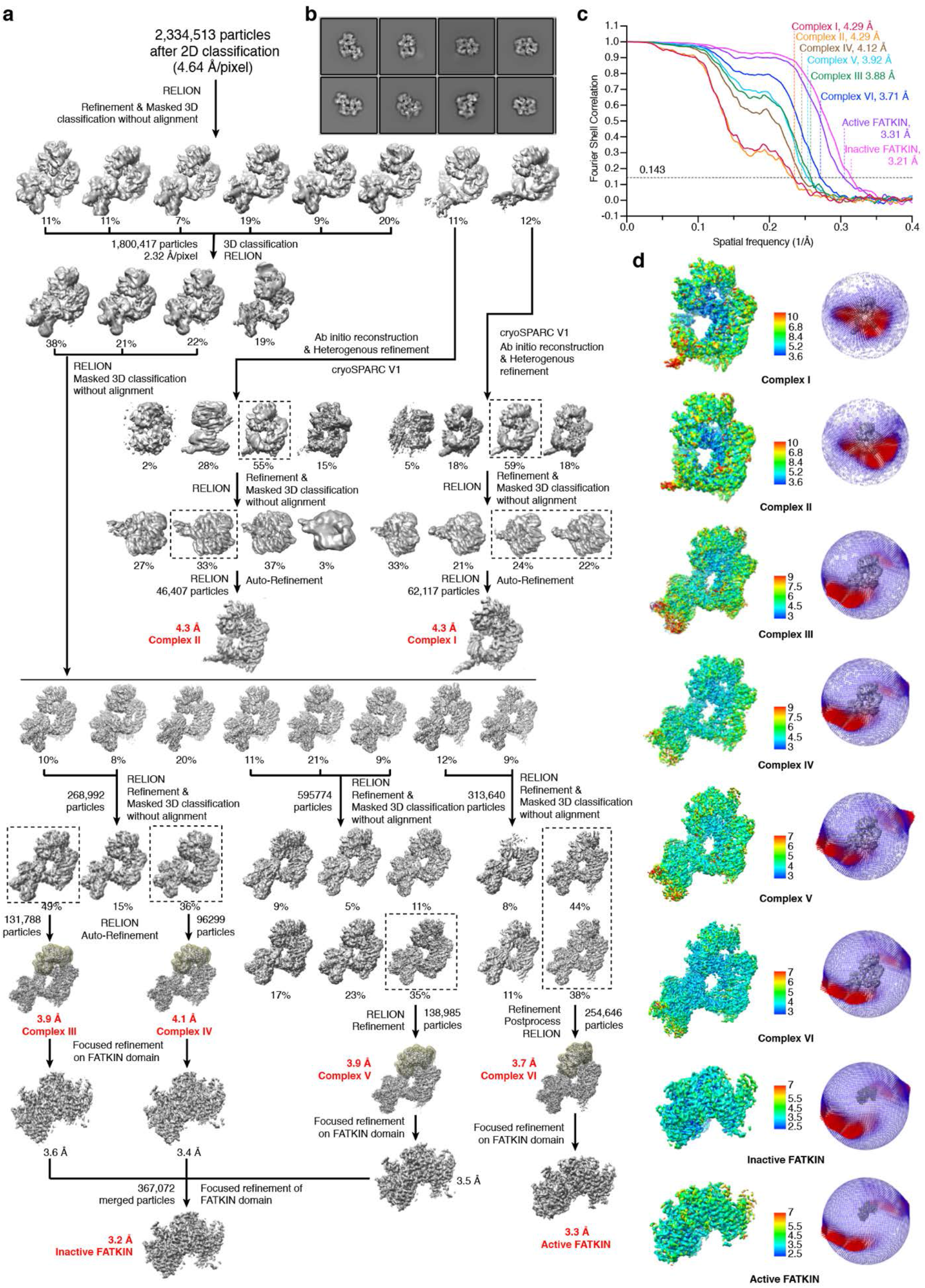
Structure determination of DNA-PK complexes by cryoEM. **S1a.** After initial 2D and 3D classification, further 3D classification with and without alignment led to the final six DNA-PK complexes and two locally refined FATKIN structures. **S1b.** Representative 2D classification results. **S1c.** FSC analysis of the quality and map resolution of each complex structure. **S1d.** For each complex, a surface presentation of its map, colored according to the local resolution estimated by ResMap and the scale bar on the side, and angular distributions of all particles used for the final three-dimensional reconstruction are shown.

**Fig. S2.**
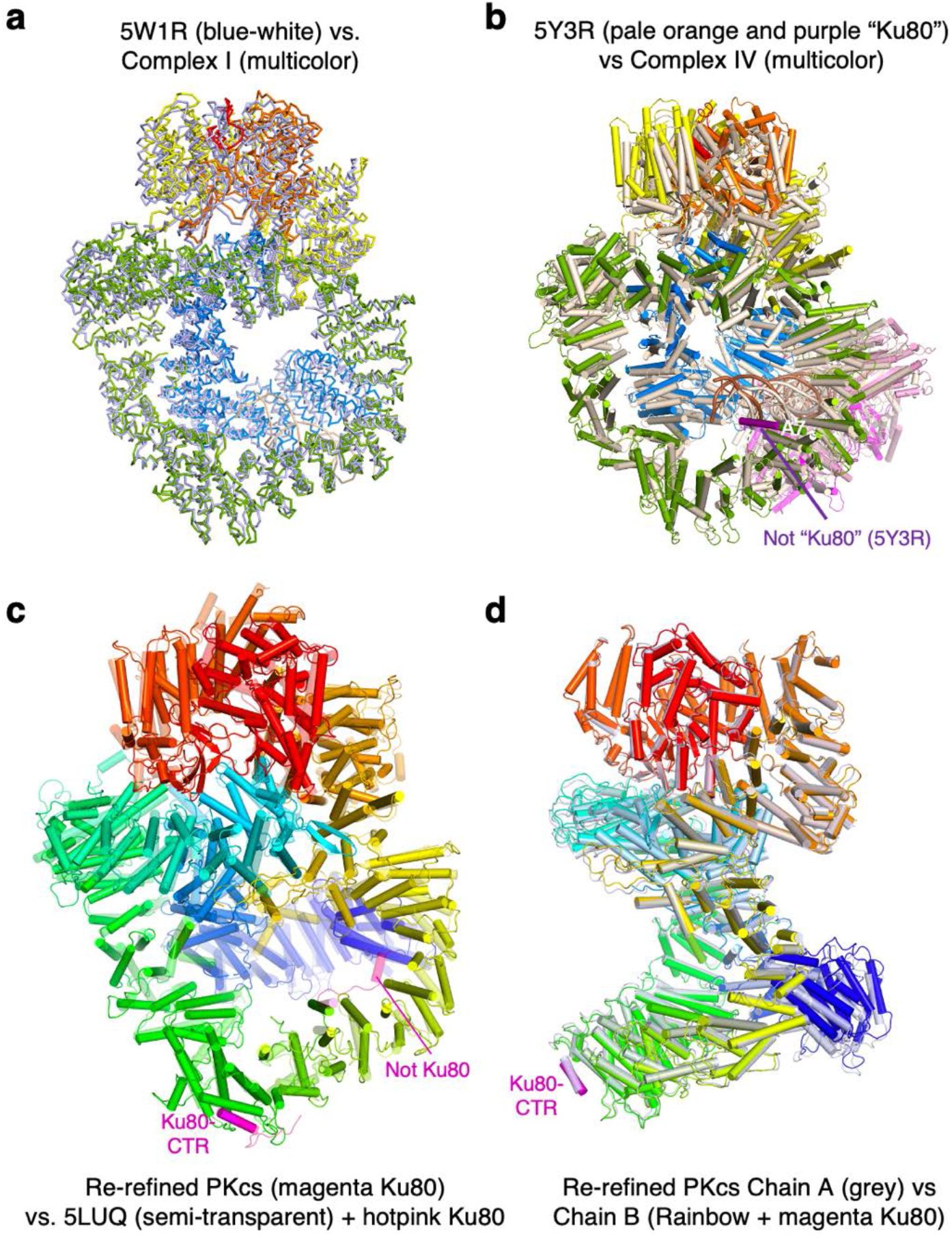
Structural comparison with previously published DNA-PK or DNA-PKcs. **S2a.** Ribbon superposition of DNA-PKcs in complex I (multicolor) and 4.4 Å cryoEM structure of DNA-PKcs alone (PDB: 5W1R, in blue-white). **S2b.** Cartoon superposition of complex IV (multicolor as in Fig. 1a) with the previously reported 6.6 Å DNA-PK complex (PDB: 5Y3R, in wheat color). The purported Ku80 helices (dark purple) turned out to belong to DNA-PKcs. **S2c.** Superposition of the re-refined (PDB: 7K17) (solid color) and the original crystal structure of DNA-PKcs (PDB: 5LUQ, semi-transparent) in rainbow colors (from blue N- to red C-terminus). The Ku80 CTR helices in the original model are colored pink. Only the very C-terminal one remains in the re-refined structure. The other two belong to DNA-PKcs. **S2d.** Two copies of DNA-PKcs in each asymmetric unit of the re-refined crystal structure are superimposed to show differences between them. One has rainbow colors and the other is colored in blue-grey.

**Fig. S3.**
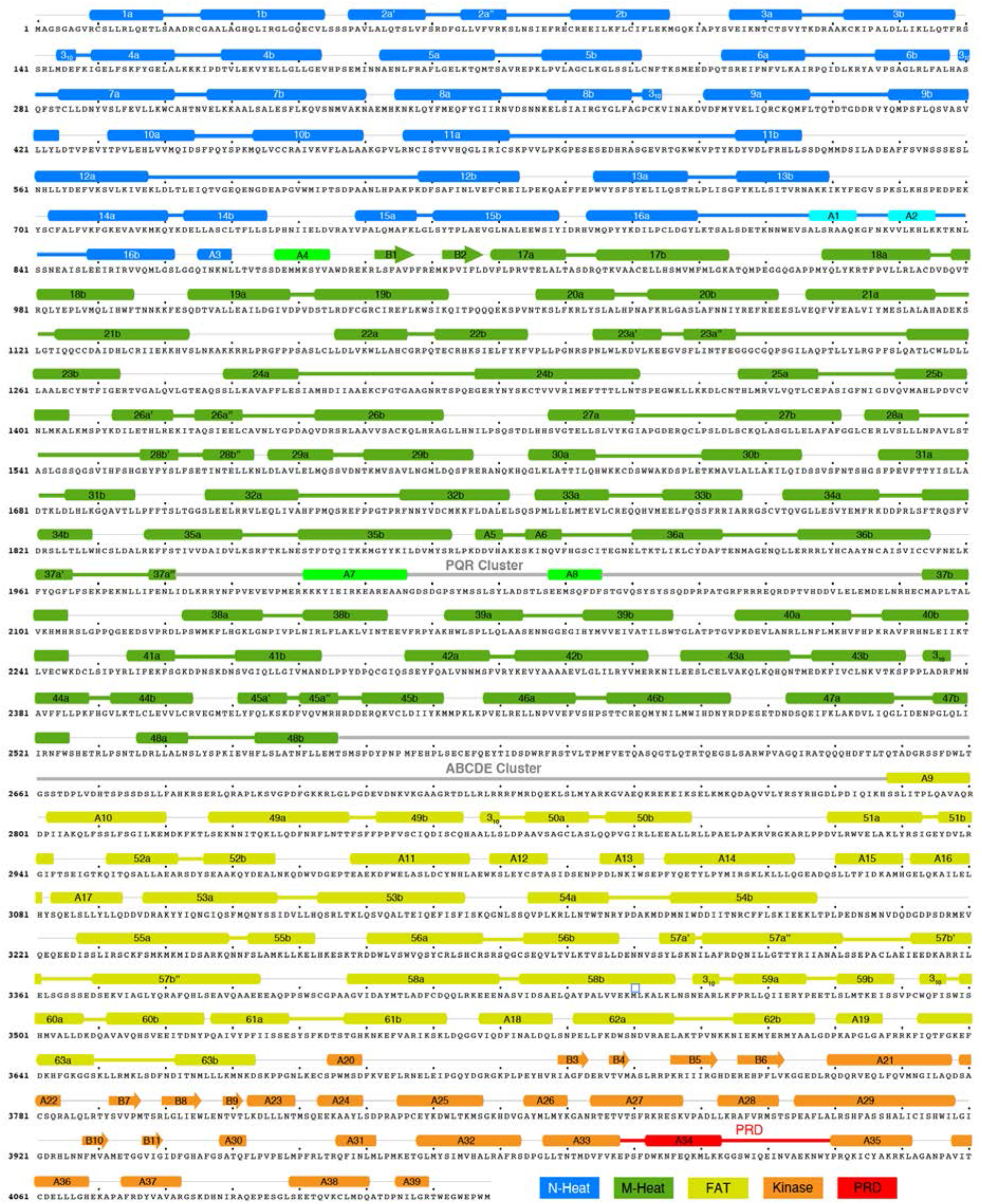
The primary and secondary structures of DNA-PKcs. Helical repeats are numbered from 1 to 63, and two helices in each repeat are labeled “a” and “b”. Non-repeat a helices and β are named sequentially as A1-A39 and B1-B11, respectively. Secondary structures are defined based on the 3.7 Å complex VI structure. ABCDE and PQR clusters of DNA-PKcs autophosphorylation sites are marked. Colorcoding scheme for domains and associated secondary structures is shown at the bottom.

**Fig. S4.**
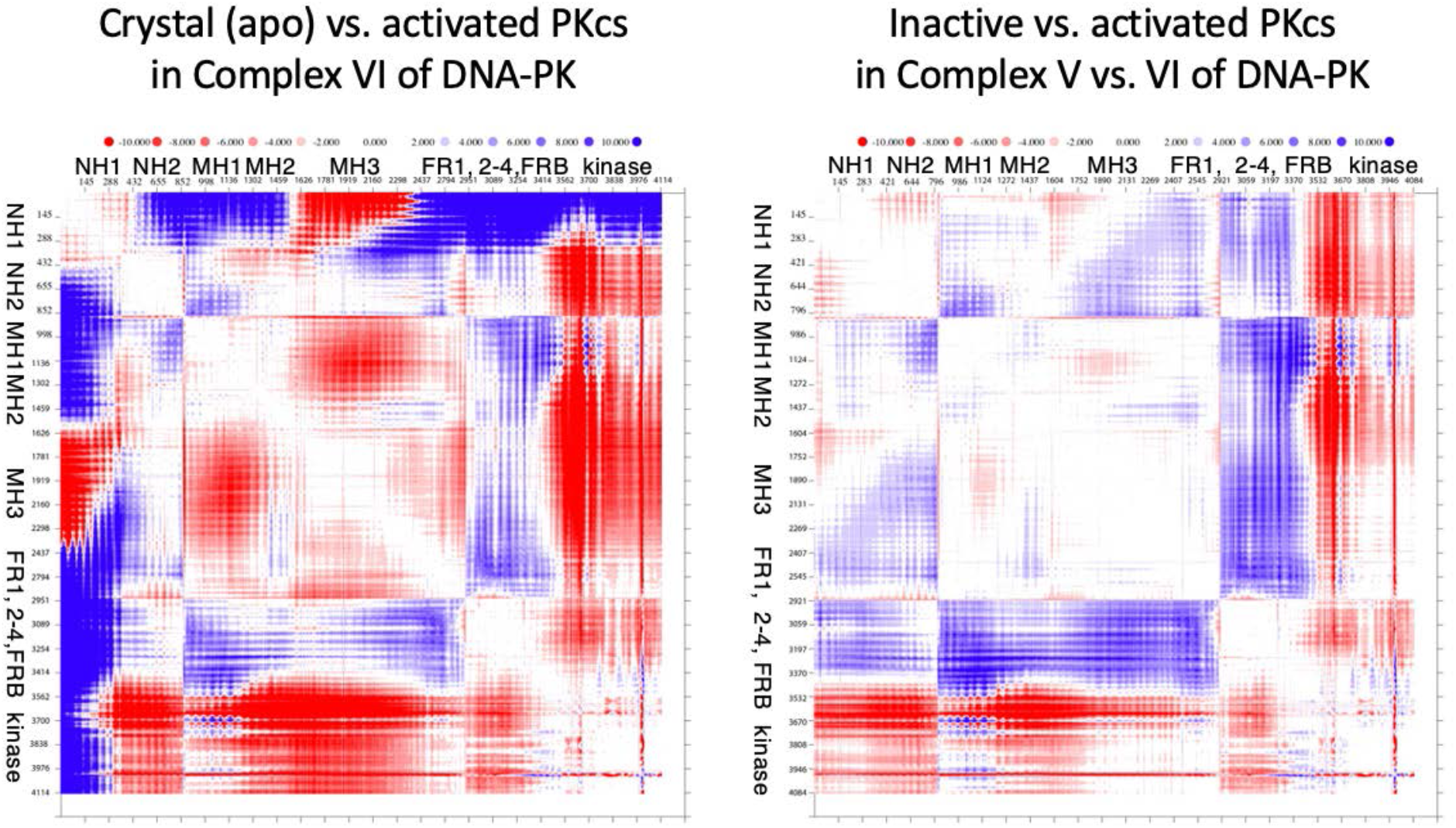
DDMP analysis of domains in DNA-PKcs. DDMP of DNA-PKcs between the re-refined crystal structure and complex VI (activated) (left) and of DNA-PKcs between complexes V and VI (right). Blue and red color indicate positive and negative distance changes. The maximal color scale is set to 10 Å.

**Fig. S5.**
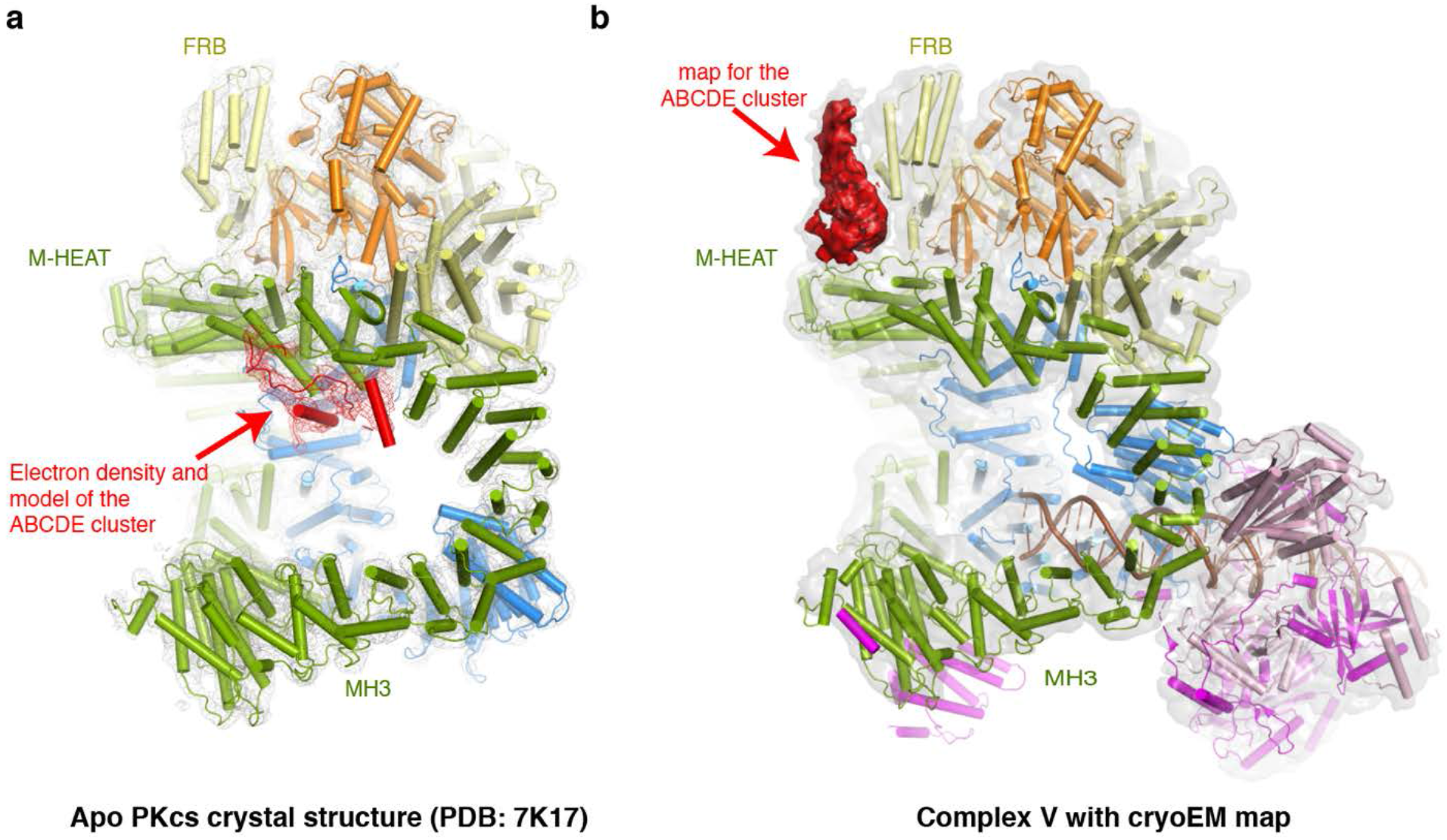
Conformational changes of ABCDE cluster in apo DNA-PKcs and DNA-PK bound to DNA. **S5a.** Cartoon model of the ABCDE cluster in chain A of re-refined crystal structure (PDB: 7K17) is superimposed with red electron density map (2Fo-Fc) contoured at 1 σ. **S5b.** The uninterpreted portion of unsharpened 3.74 Å cryoEM map in DNA-PK complex V, which belongs to the ABCDE cluster, is contoured at 5 σ and shown in red.

